# Treatment of Mixed Azo Dyes in an Aerobic Sequential Batch Reactor and Toxicity Assessment using *Vigna radiata*

**DOI:** 10.1101/680249

**Authors:** Akshaya Vidhya T, K Veena Gayathri, Tasneem M Kathawala

## Abstract

Azo dyes are the most widely used dyes in the textile industry due to their stability buttheir redundancy to degradation is of major concern, particularly to aquatic ecosystems.Unbound dye is let out in the effluent which not only adds to pollution but its toxic metabolites are known to be carcinogenic leading to severe cases of disease. Biological degradation and toxicity removal has been shown to be an easy and effective process for treating textile effluents. In the present study, a laboratory scale aerobic sequential batch reactor (SBR) was designed and operated for the analysis of degradation of mixed reactive azo dyes. Biological degradation was carried out by activated sludge process at an alkaline pH (8.5). Reactive Brown, Reactive Black and Reactive Red dyes were used in the study at a concentration of 100, 500 and 1000 mg/L in synthetic waste water. The effect of increasing dye concentration on the decolorization efficiency, COD and BOD removal along with chloride, hardness, TDS, MLSS and MLVSS was monitored. The COD removal increased from 34% to 61.15% and then dropped to 21.16% at the highest used concentration. The BOD removal decreased from 63% to 55.55% to 28.14% with increasing dye concentration. In order to remove the residual dye from the effluent, a biosorption experiment was also conducted using dried activated sludge (DAS). The DAS successfully removed more than 0.300 mg of dyes by absorption within 2 hours. A toxicity assessment was carried out by mean of a phytotoxicity test on *Vigna radiate* where the percentage of germination was used to detect toxic effects of untreated dye containing wastewater on plant growth. The treated wastewater showed 100% germination compared to 70% in untreated wastewater containing 100 mg/L mixed dyes confirming the efficacy of the treatment.

## Introduction

Synthetic dyes used most frequently in textile industries are major contributors to causes of pollution. The participation of textile industries as polluters of water bodies has witnessed an alarming increase in the past few decades [1]. Playing the lead role, are Azo dyes which are the most commonly used dyes and possess the characteristic azo (–N=N-) group. These highly stable dyes find their applications in several other industries apart from the textiles, like cosmetics, paper, food and leather [2]. The colour affects the penetration of sunlight and thus aquatic plants [3] and the high nitrogen content often leads to eutrophication [4]. The consequences associated with an inefficient treatment of textile wastewater involve several risks. More than 25% of amine based dyes are identified as carcinogens and several being carcinogenic to humans [5]. Hence effluents containing these xenobiotic and recalcitrant dyes cannot be degraded by conventional wastewater treatment methods [6] and this makes it imperative to develop a suitable and efficient method to contain the pollution caused by these dyes.

The conventional methods of treating wastewater include physical, chemical and biological methods. Physical and chemical treatments are not efficient in degrading Azo dye contaminated effluent and they also produce secondary waste products that would require further treatment [7]. Biological treatment involves several microorganisms that accumulate and/or degrade dyes and chemicals present in wastewater [8]. This natural and environmental friendly method is surprisingly effective and has several advantages over the other methods. It is cost effective and the number of secondary pollutants produced is far lesser in comparison to other treatment methods [9].

The biological treatment system includes the activated sludge process (ASP) which is a widely used procedure in wastewater treatment. The ASP can be carried out by aerobic, facultative and anaerobic microorganisms under appropriate oxic/anoxic conditions. The Azo dye is degraded by a consortium when it acts as an electron acceptor, causing the reduction of the azo bond and resulting in the formation of aromatic amines under anaerobic conditions [10]. The aerobic process involves degradation of pollutants by flocculating biomass in an aeration tank.

The sequencing batch reactor (SBR) gives promising results for the degradation of dyes. The sequencing batch reactor operates in time and it is fill and draw type of activated sludge process. An SBR can perform equalization, neutralization, biological treatment and secondary clarification in a single tank [11]. There are numerous studies done to find a suitable process for reusing the textile wastewater. For the possibility of reusing the textile effluent, bioreactor studies are done [12]. In biological treatment methods, bioreactors are used for effective biomass production that can degrade the dyes present in wastewater.

The primary objective of the present study is to develop a suitable reactor system for efficient degradation of mixed dye. A batch study was performed to evaluate the degradative efficiency of the microbial consortium present in the sludge. The study was proposed to treat synthetic textile effluent (containing Reactive mixed dyes i.e. Reactive Red, Reactive Black and Reactive Brown) using Sequencing Batch Reactor. Their operational parameters such as COD, BOD, hardness, chloride, TDS, pH and temperature were studied through the scale up study.

## Materials and methods

### Preparation of synthetic wastewater

The composition of synthetic wastewater containing dipotassium hydrogen phosphate (0.5 g/L), potassium dihydrogen phosphate (0.5 g/L), magnesium sulphate (0.1 g/L), ferric chloride (0.025 g/L), ammonium chloride (1 g/L), calcium chloride (0.01 g/L), yeast extract (0.025 g/L) and lactose (0.15 g/L) [13]. The pH was adjusted to 8.5 and autoclaved. The dyes were membrane filtered and added to the prepared synthetic wastewater after autoclaving. Each dye Reactive Red (RR), Reactive Brown (RBr) and Reactive Black (RB) was added in different concentration that is 100 mg/L, 500 mg/L and 1000 mg/L. The dyes were purchased from India Mart. The chemicals used for the synthetic wastewater were purchased from Merck.

### Batch studies for decolorization of mixed dyes

To check the efficiency of dye removal in the synthetic wastewater by the activated sludge, batch studies were performed. In a 200 ml conical flask 100 ml of synthetic wastewater along with the three mixed dyes and 100 ml of collected activated sludge was mixed and kept in orbital shaker and incubated at 30° C for five days. A control was also kept with distilled water instead of activated sludge and kept for incubation in an orbital shaker. The samples were collected everyday from the conical flask, centrifuged at 8000 rpm for 10 minutes and diluted and decolorization percentage was determined using a UV-Visible spectrophotometer at wavelengths of 369 nm for mixed dyes, 518 nm for RR, 440 nm for RBr and 614 nm for RB. A graph was plotted to determine the decolorization percentage.

### Operational setup

Activated sludge was collected at five days interval from the aeration tank of Koyambedu Sewage Treatment Plant (STP), Chennai. For the activated sludge process, a 14 L bioreactor was built with a height of 30 cm and a breath of 24 cm, made up of acrylic tank. For aeration, a porous air diffuser was attached to the bottom and for homologous mixing, a stirrer was used at 150 rpm. Each cycle of SBR had five periods they are FILL (30 minutes), REACT (22 hours), SETTLE (1 hour), DRAW (30 minutes) and IDLE (30 minutes). Throughout the process the temperature was maintained at room temperature (37°C). A working volume of 8 L in which 4 L was activated sludge and 4 L was synthetic wastewater with the respective dye concentrations of 100 mg/L, 500 mg/L and 1000 mg/L respectively. The reactor was operated dye free for three days for the acclimatization of the sludge to the synthetic wastewater. The effluent from the SBR was collected during the DRAW phase and chemical analyses were done using the standard methods. The degradation of reactive red, reactive black and reactive brown was determined using a UV-Vis spectrophotometer at the respective nm. After completion of every concentration of the synthetic dyes, the concentrations were further increased. The COD removal, BOD removal, MLSS and MLVSS were determined regularly. Figure 1 depicts the operational setup of the sequencing batch reactor.

**Fig 1.**
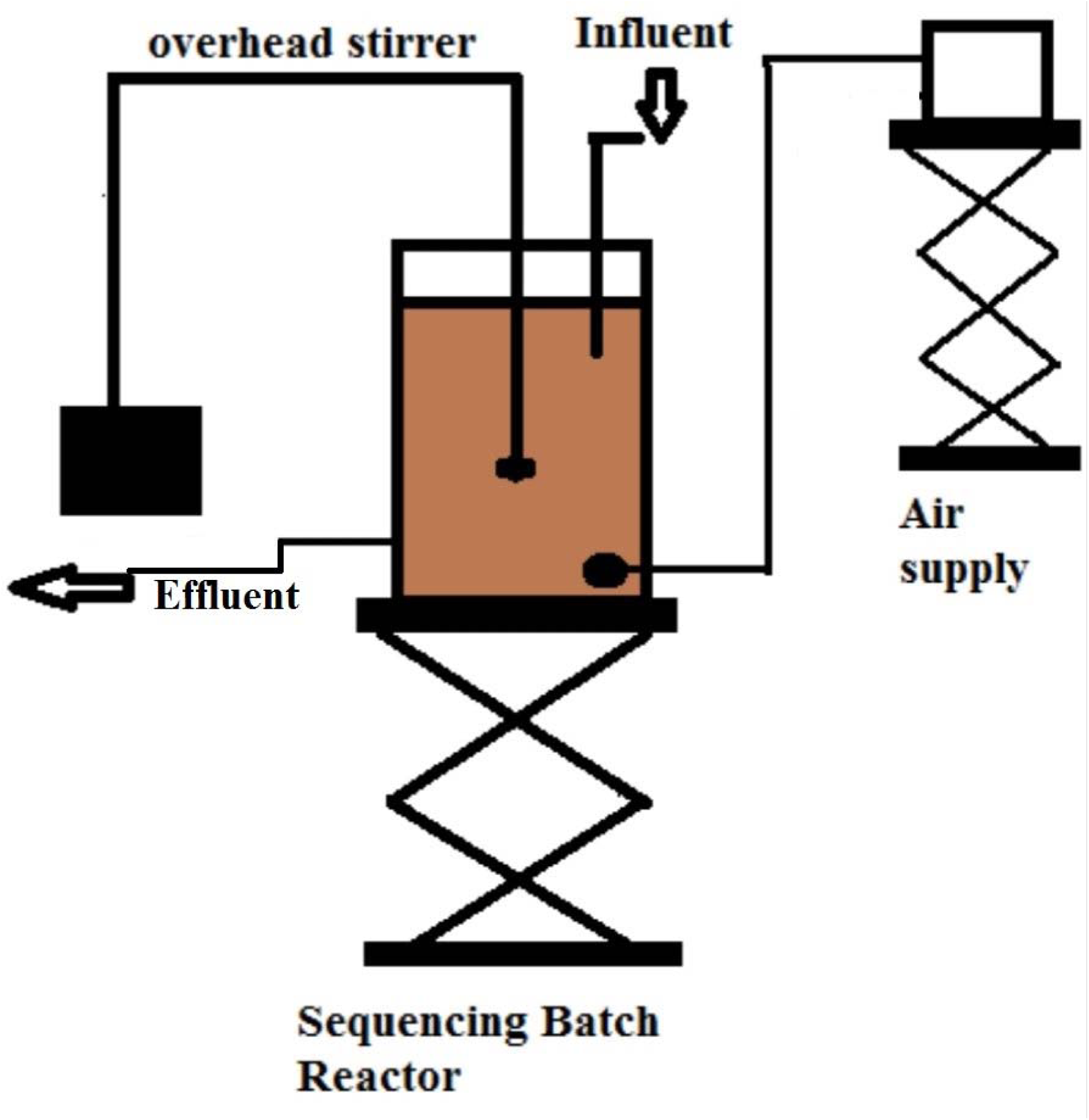
SBR setup for treatment of wastewater.

### Analyses

For the analysis of physiochemical parameters, standard methods were followed for the determination of COD, BOD, TDS, hardness and chloride [14]. The pH was checked during the treatment with a pH meter and was maintained at 8.5. The temperature changes were measured using a thermometer.

### HPLC Analysis

The wastewater treated from the reactor containing the degraded dyes were filtered in 0.45μm membrane filter. The filtrates were then extracted with equal amount of ethyl acetate and flash evaporated in rotary vacuum evaporator in temperature controlled water bath (50°C) and residues were dissolved in 2 ml of HPLC grade methanol and used for HPLC analysis. The HPLC model was Shimadzu Prominence Binary Gradient HPLC System. The extracted samples were analyzed by HPLC having a mobile phase of 100% methanol [15].

### Biosorption studies on treated wastewater

A batch study using dried activated sludge was carried out in order to obtain further decolorization of the SBR effluent. Activated sludge obtained from the aeration tank of Nesavakam sewage treatment plant, Chennai was centrifuged and the pellet was washed thoroughly using distilled water. The sludge was spread in a sterile petriplate and was kept in hot air oven at 60°C overnight. The dried activated sludge (DAS) was scraped out from the petriplate gently and grinded to fine powder using mortor and pestle. It was then seived through 105 μm diameter mesh and stored in a dry bottle and was used as biosorbent for the experiment [16].

The study was carried out in 250 ml Erlenmeyer flasks in which 100 ml of the 100 mg/L treated wastewater was taken and 0.5 g of the dried activated sludge was added to it. The initial absorbance was checked using a UV-Visible spectrophotometer and was kept in orbital shaker. Samples were collected at 20, 40, 60, 80, 100 and 120 minutes, centrifuged at 10,000 rpm for 10 minutes and the absorbance was determined in the spectrophotometrically. The obtained values were substituted in the following formula to find the amount of dye that was adsorbed by the biomass. A control was maintained with 100 ml of treated wastewater without dried activated sludge.

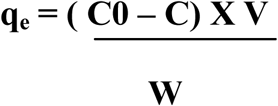

where,

**q**_**e**_ is the amount of dye adsorbed by the biomass

**C**□ is initial concentration of dye

**C** is final concentration of dye

**V** is volume of dye solution

**W** is weight of the biomass

### Phytotoxicity study

The phytotoxicity study was carried out with *Vigna radiata* (green gram) at room temperature. Three different pot containing 20 seeds of green gram in each were watered with 10 ml of distilled water as control, untreated synthetic wastewater and treated synthetic wastewater respectively [16]. Germination percentage was calculated by using the below formula

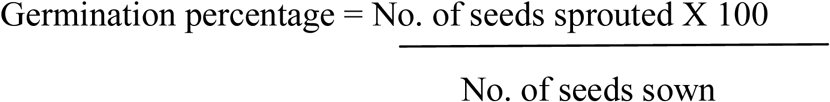

## Results and Discussions

### Decolorization of mixed dyes

Batch studies were conducted to study the removal of mixed reactive dye by the microbes present in activated sludge. It was observed that the mixed reactive dyes removal was estimated up to 40 % of mixed dye, 39 % of RR, 56 % of RB and 42 % of RBR respectively in aerobic condition under room temperature and at a pH of 8.5 at 150 rpm in five days. Figure 2 shows the decolorization of the mixed azo dyes in a five day batch study.

**Fig 2.**
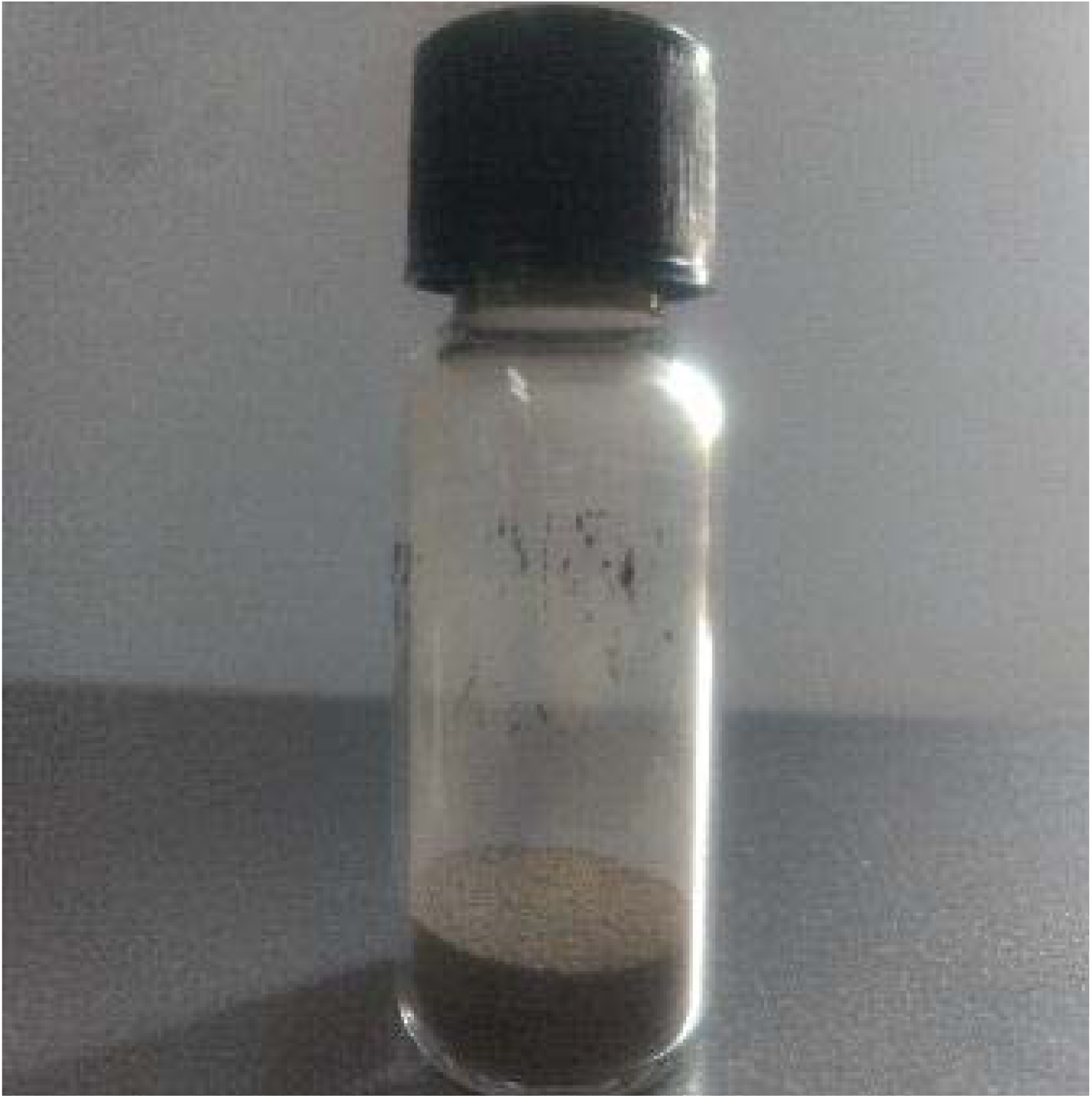
Dried activated sludge prepared for biosorption studies.

### Sequential Batch Reactor system with 100mg/L of mixed reactive dyes

Initially the SBR system was loaded with 100 mg/L of the mixed reactive dyes. The pH was maintained at 8.5. The initial concentration of COD in the synthetic wastewater was 320 mg/L on the first day of seeding which was reduced to 110 mg/L after 10 days of SBR operation. The COD of synthetic wastewater was half the concentration of about 160 mg/L on second day after which the treatment showed active degradation of mixed dyes present in synthetic wastewater by the microbial community. The COD reduction was observed in the graph which gradually decreased till the tenth day. After tenth day the amount of bacteria in the activated sludge couldn’t reduce COD and it remained the same. At the end of the operation the COD was 110 mg/L therefore 34.375 % of COD was removed in ten days. Tenth day was permissible level for the bacterial consortium present in the activated sludge to reduce COD levels.

Utilization of mixed dyes was also indicated by BOD levels in the wastewater. Initially BOD concentration of wastewater was about 900 mg/L, only one fourth of it was reduced to 570 mg/L on second day. There was an increase in BOD to 660 mg/L on the third day and 810 mg/L on the fifth day showing the competition between the microbes to survive in the wastewater while the percentage of BOD drastically decreased to 480 mg/L on tenth day and showed 63 % of BOD removal.

The total dissolved salts (TDS) were 600 mg/L initially which gradually decreased to 400 mg/L. There was a sudden drop of TDS during day two which again increased to 700 mg/L. There was no decrease in TDS after day six. The chloride content in the synthetic wastewater was 400 mg/L, which was 350 mg/L end of the treatment in SBR.

Mixed Liquor Suspended Solids and Mixed Liquor Volatile Suspended Solids were initially at 2000 mg/L and 1000 mg/L which increased to 6000 mg/L and 3000 mg/L at the end of the process showing the increase in the microbial population in the activated sludge by utilizing the dyes as carbon source in the wastewater. There was a drastic increase during day four to six during which the MLSS and MLVSS was 8700 mg/L and 4400 mg/L which proved the growth of microorganisms were high during day four to six. After which the population decreased at the end of the studies which may be due to decrease in dye concentration which was utilized by the microbes present in the activated sludge.

The hardness of water was observed to be 700 mg/L which is considered to be showing highest hardness. After the treatment it was observed that 700 mg/L of hardness decreased to 300 mg/L of hardness on the tenth day.

### Sequential Batch Reactor system with 500mg/L of mixed reactive dyes

Initially the SBR was loaded with 4000 mg/L of COD, which was removed by the microbes present in the activated sludge to 1554 mg/L at the end of the SBR operation. The COD decreased to 2400 mg/L on third day but it increased again to 3440 mg/L on fifth day showing the concentration of dye is high for the microbes in the activated sludge to degrade. On the sixth day from 3440 mg/L, the COD decreased to half the concentration to 1600 mg/L by which it could be observed that the activated sludge got acclimatized to the wastewater with 500 mg/L dye concentration and therefore they were able to decrease the COD level. There was an increase in COD concentration on day eight where the COD was 2640 mg/L which decreased on ninth and tenth day to 1900 mg/L and 1554 mg/L respectively. Therefore during the end of the operation in SBR, 61.15% of COD removal was achieved. The system was able to maintain the COD removal at 61.15 % showing that the microbes were able to degrade the dyes and other recalcitrant substance in the seeded wastewater.

The MLSS and MLVSS was initially 1000 mg/L and 500 mg/L in the sludge that was seeded to the SBR. There was an increase in MLSS and MLVSS on second day which was 2700 mg/L and 1300 mg/L. It gradually increased proving that the microorganisms got acclimatized therefore the population of microbes in the reactor increased till seventh day which was 7400 mg/L and 3500 mg/L, after which the microbes lost their ability to adsorb the dye in the synthetic wastewater which is the carbon source. Therefore there was no increase in population after seventh day. During the end of SBR operation the MLSS and MLVSS were 6600 mg/L and 3300 mg/L.

The synthetic wastewater that was used in 500 mg/L dye concentration studies contained 900 mg/L of BOD initially. On the second day the BOD increased to 1290 mg/L and it again increased to 1620 mg/L on third day by which we can observe that the dye concentration was too high for the microorganisms in the reactor. They were not able to decrease the BOD level till third day, after which the BOD decreased to 600 mg/L showing how the microbes in the activated sludge trying to acclimatize to the high concentration of dye. The BOD level increased to 780 mg/L on fifth day after which there was no increase in BOD level but gradual decrease till the end of the operation by which we can observe that the bacteria in sludge got acclimatized to 500 mg/L dye concentration. On tenth day, BOD was 400 mg/L, therefore the total percentage of BOD removal observed in SBR was 55.55 %.

TDS was 900 mg/L initially and was decreased to 650 mg/L. Therefore 27 % of TDS was decreased in SBR.

The hardness for water on the first day was 1000 mg/L which was considered to be of highest hardness. The hardness of water increased to 1500 mg/L. After fourth day the hardness of water gradually decreased to 400 mg/L at the end of the operation.

The chloride content in the synthetic wastewater that was seeded to the reactor was 1300 mg/L which decreased gradually to 760 mg/L in ten days of SBR operation. Therefore 41.53 % of chloride was removed by the acclimatized microbes in the sludge.

### Sequential Batch Reactor system with 1000mg/L of mixed reactive dyes

In 1000 mg/L studies, the pH fluctuated from 8.5 to 8, it was maintained at 8 till the end of the operation and the temperature was room temperature.

The influent for 1000 mg/L studies contained high concentration of 6000 mg/L of COD. There was a gradual decrease in COD concentration everyday but since it was 1000 mg/L studies, the microorganisms couldn’t get adapted to the high concentration of COD and there was only a mild reduction in COD, which gradually decreased to 5742 mg/L on sixth day after which it reduced rapidly to 4700 mg/L and then increased on eighth day to 4770 mg/L, on ninth day it reduced to 4730 mg/L and it remained constant for few more days by which we can understand that the microorganism lost their ability to reduce COD. The amount of COD removed in ten days of SBR operation was 21.16 %.

The BOD concentration was 1670 mg/L in the influent which reduced gradually to 1200 mg/L in the end of the process by which we can observe that the bacteria in activated sludge tried acclimatizing to the synthetic wastewater with high concentration of dye, after tenth day they lost the ability to degrade the dyes and the BOD and COD remained constant. The amount of BOD reduced was 28.14 %.

TDS was 900 mg/L, it reduced to 700 mg/L on day five and remained constant till day seven after which it reduced to 654 mg/L and reached 600 mg/L on day ten. In ten days 33.33 % of TDS removal was achieved.

The MLSS and MLVSS on the day of seeding was 3000 mg/L and 1500 mg/L. Since the concentration of dye was high in 1000 ppm studies, the sludge that was seeded to the SBR contained high population of microorganisms for efficient degradation. The population remained constant for two days after which the microorganisms slowly acclimatized to the synthetic wastewater and it increased to 5300 mg/L of MLSS and 2600 mg/L of MLVSS on sixth day. Once the carbon source in the synthetic wastewater decreased after sixth day, the population of microorganisms started to decrease and reached 4000 mg/L of MLSS and 2000 mg/L of MLVSS at the end of the operation in SBR.

Initially chloride concentration was 1700 mg/L which decreased to 1400 mg/L on third day. There was rapid decrease after third day, it reached 1016 mg/L on fourth day. On tenth day it was 833 mg/L. Chloride removal in ten days of operation was 51 %. Since it was 1000 mg/L, the hardness of water was extremely high i.e., 1400 mg/L, which reduced to 900 mg/L on fourth day and 551 mg/L on day ten, 60 % of hardness reduction was achieved in SBR.

The results obtained at concentration of 100mg/L were comparable to those obatained by Shaw et al (2002) where COD removal was found to occur between 70-80%. Tantak and Chaudhari (2006) reported 81.95, 85.57 and 77.83% of COD in aerobic SBR following Fentons’ oxidation. El-Gohary and Tawfik (2009) reported 78% removal of COD and 68.9% removal of BOD in aerobic SBR at h HRT following a coagulation flocculation. Hence, chemical pre treatments can be performed to increase the efficiency of COD, colour and BOD removal in an aerobic sequential batch reactor.

### HPLC analysis

A HPLC system, operated with UV-Vis at 369 nm was used to analyze the degradation of mixed reactive dyes in the synthetic wastewater and their intermediate. Several degraded products were also observed by HPLC analysis. By the comparison of the retention time of the initial day wastewater sample peak and the peaks of wastewater sample after treatment showed degraded products.

In the elution profile of the initial day wastewater with 100 mg/L of mixed dyes showed a high peak at 2.625 minutes and other peaks at 3.093, 3.998 and 4.614 minutes shown in figure 5 (i). After treatment in SBR for 10 days, the elution profile of SBR treated wastewater showed a reduced peak at 2.814 minutes and 3.312 103 and 4.354 minutes by which it was observed that the peak at 2.652 minutes observed in the initial day elution profile was reduced and had a shift to 2.814 minutes seen in figure 5 (ii).

**Fig 3.**
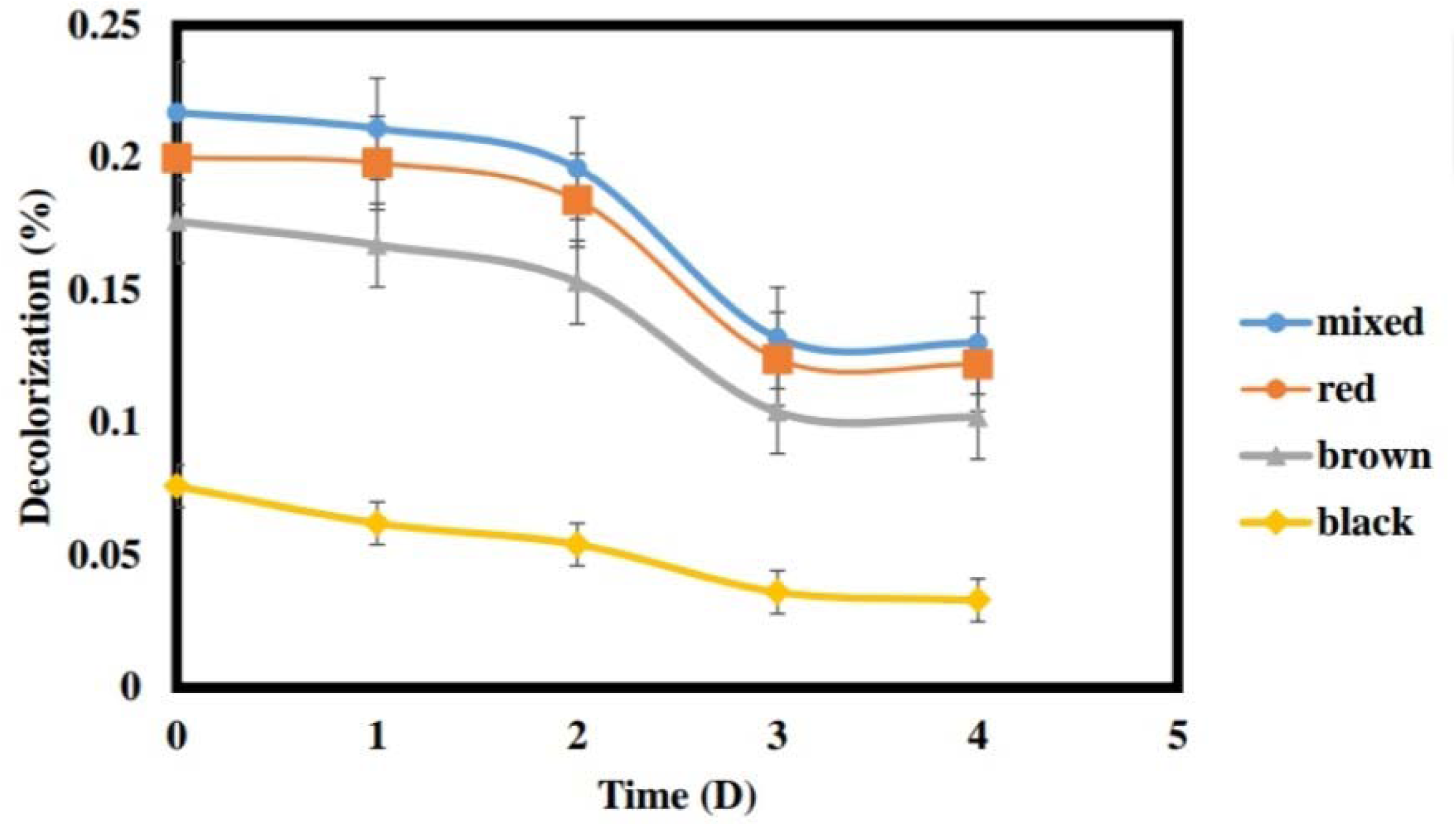
Decolorization of mixed reactive dyes in batch study.

**Fig 4.**
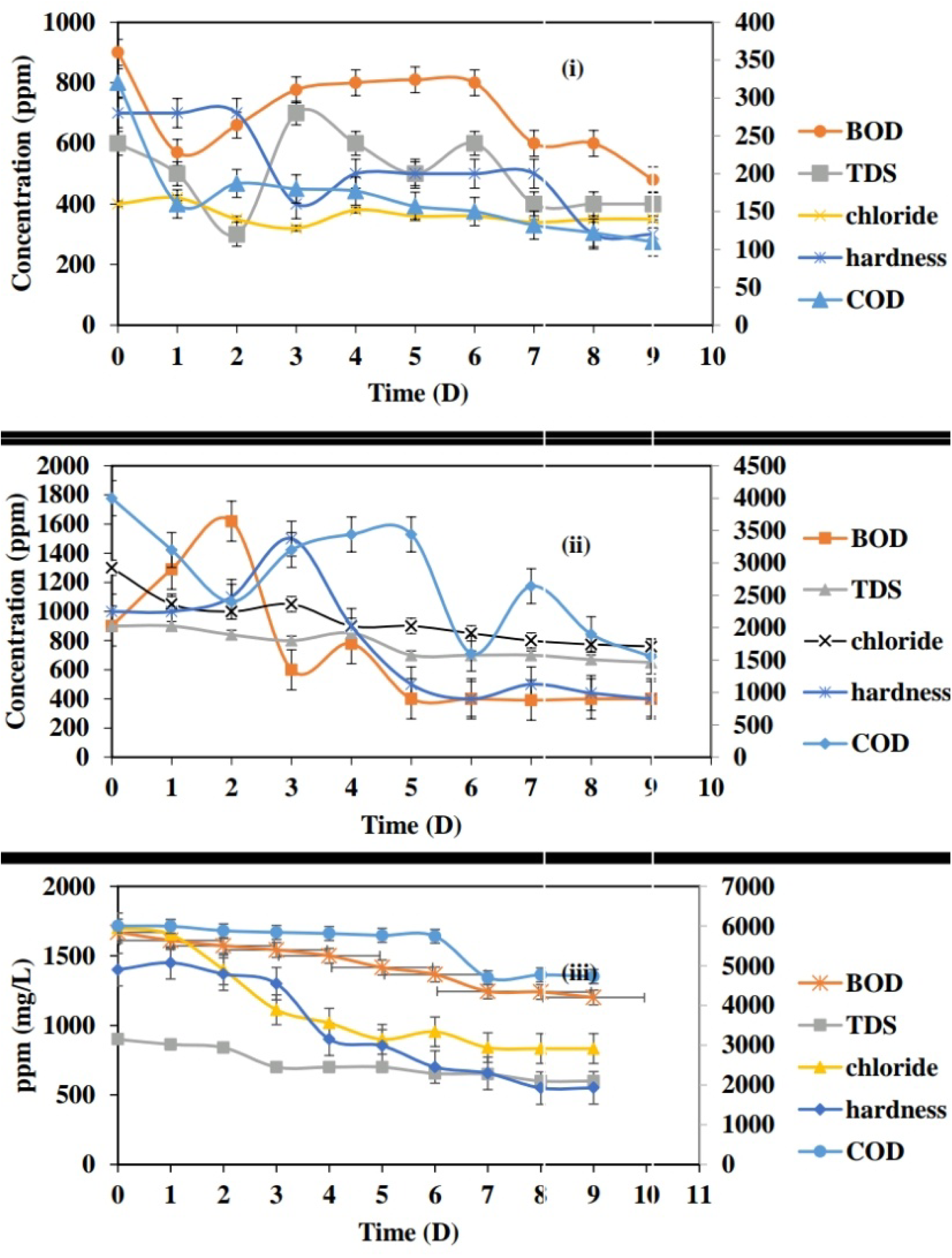
Physicochemical analyses for synthetic wastewater in SBR containing (i)100 mg/L (ii)500mg/L and (iii)1000 mg/L.

**Fig 5.**
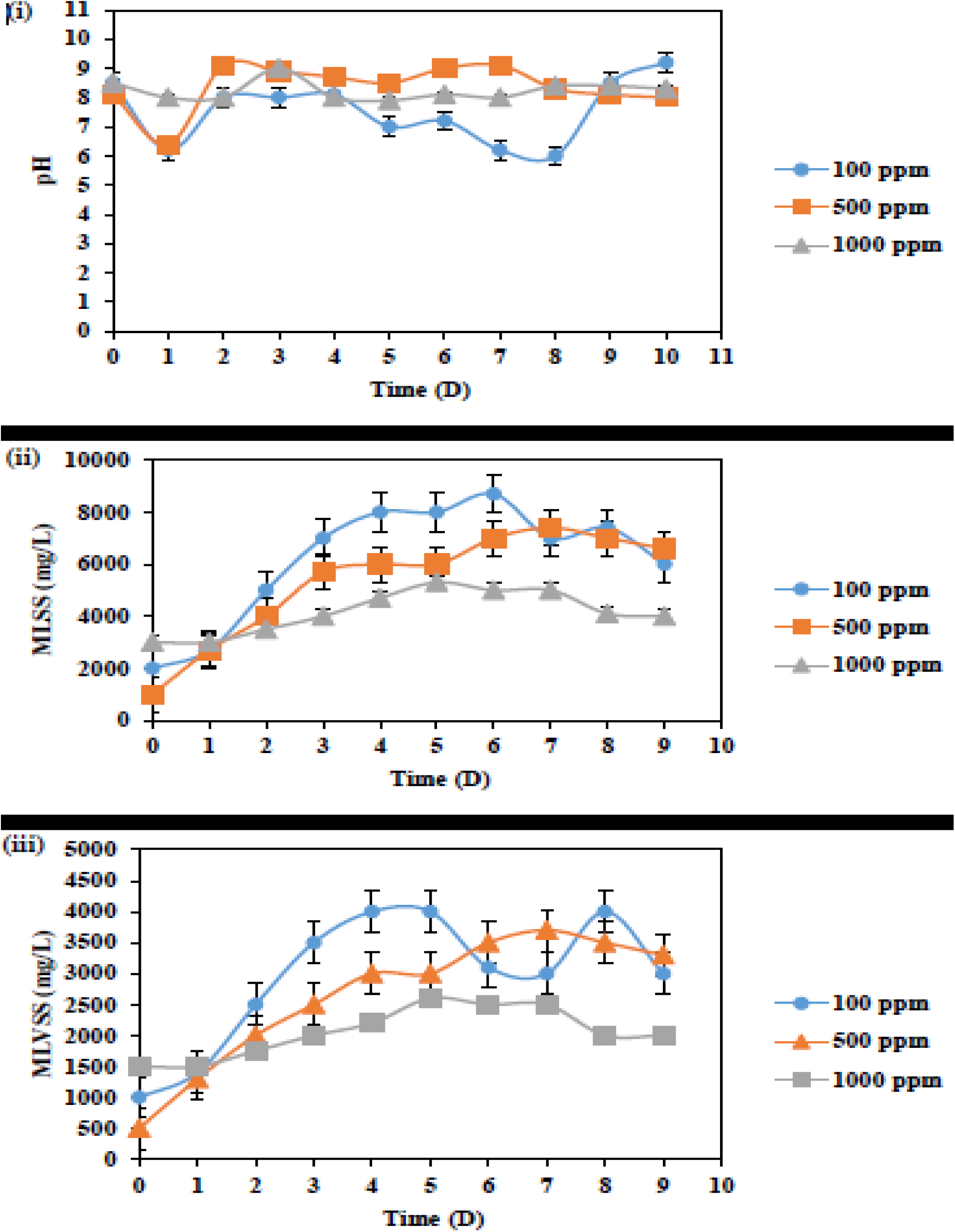
The pH, MLSS and MLVSS for synthetic wastewater in SBR containing (i)100 mg/L (ii)500mg/L and (iii)1000 mg/L.

For 500 mg/L concentration, peaks were observed at 2.638 and 3.296 retention time in the elution profile of the SBR influent shown in figure 6 (i). After the treatment of wastewater in the SBR, there was a shift in retention time to 3.252 and 4.081 and the height of the peaks were reduced which indicated that few molecules have been degraded during the treatment as seen in figure 6 (ii).

**Fig 6.**
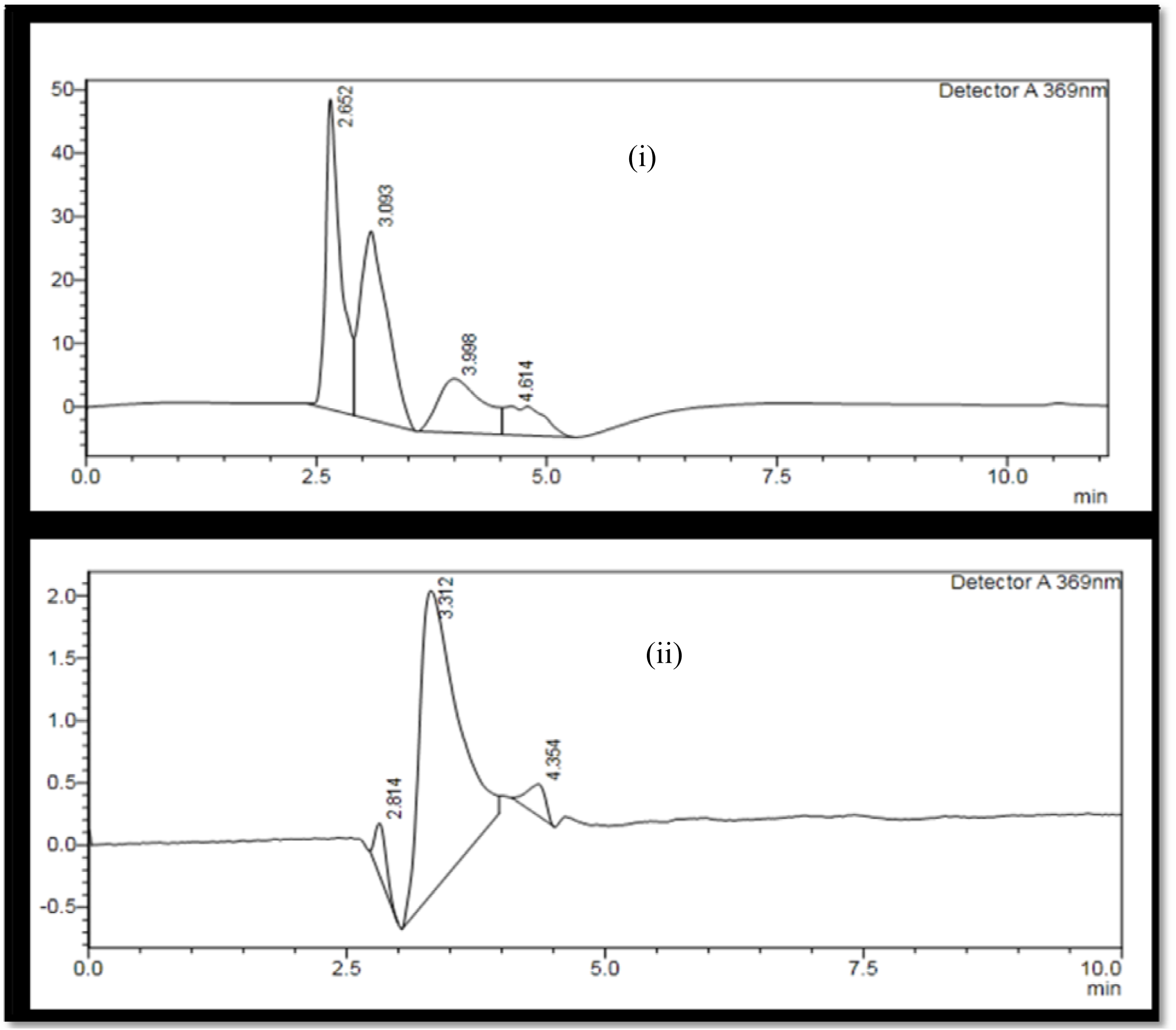
HPLC analysis of influent (i) and effluent (ii) of SBR operated with 100 mg/L of mixed dyes.

In 1000 mg/L studies, elution profile of the untreated wastewater i.e. the influent to SBR showed peaks at 1.850, 2.646, 32.896, 3.295 and 4.120 minutes of retention time shown in the below figure 7 (i). After treatment in SBR, the wastewater sample showed a shift in retention time and heights of peaks were reduced. Peaks were observed at 2.644 and 4.887 retention time by which we can observe that degradation has occurred during the treatment in SBR, shown in below figure 7 (ii).

**Fig 7.**
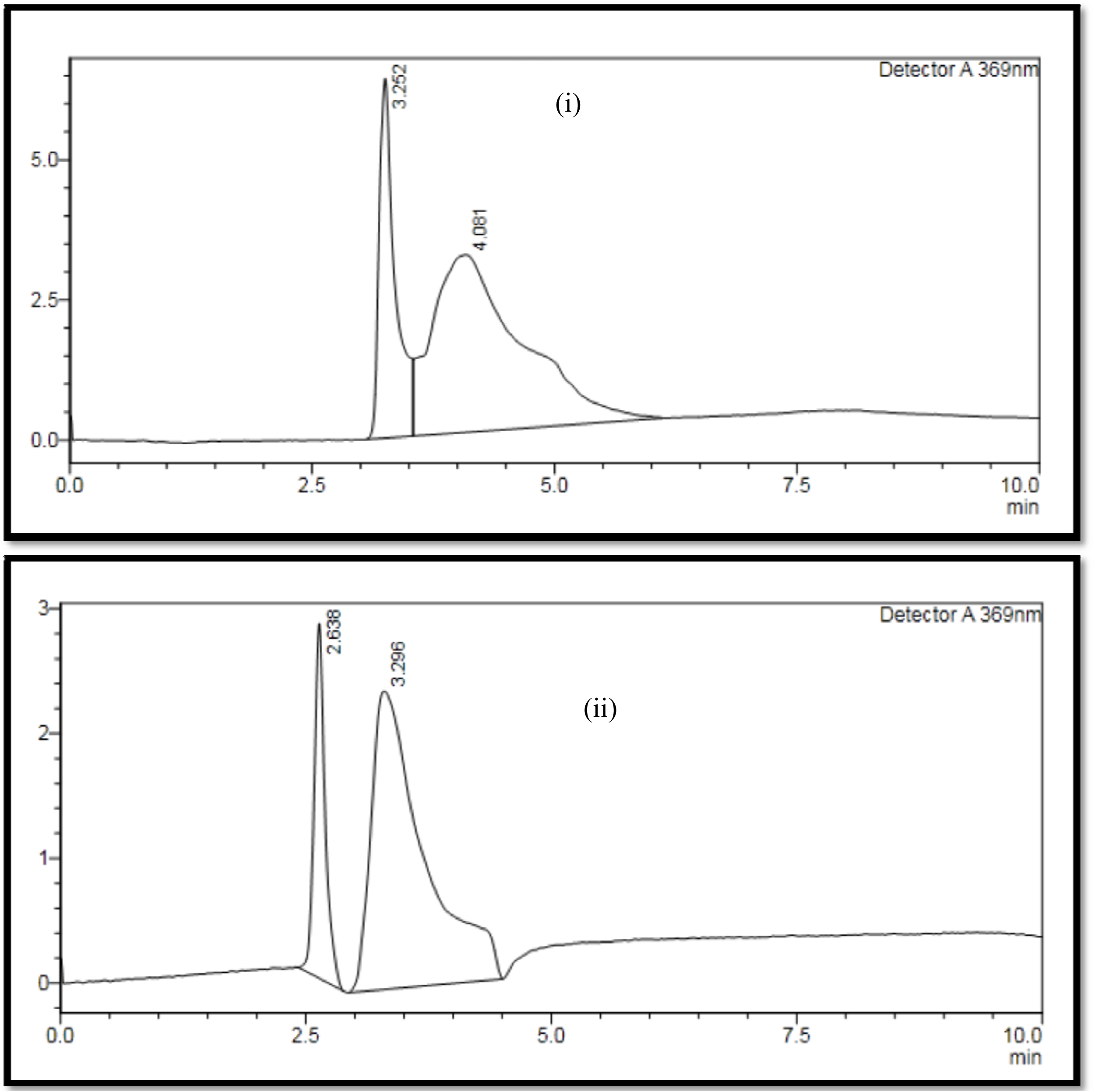
HPLC analysis of influent (i) and effluent (ii) of SBR operated with 500 mg/L of mixed dyes.

**Fig 8.**
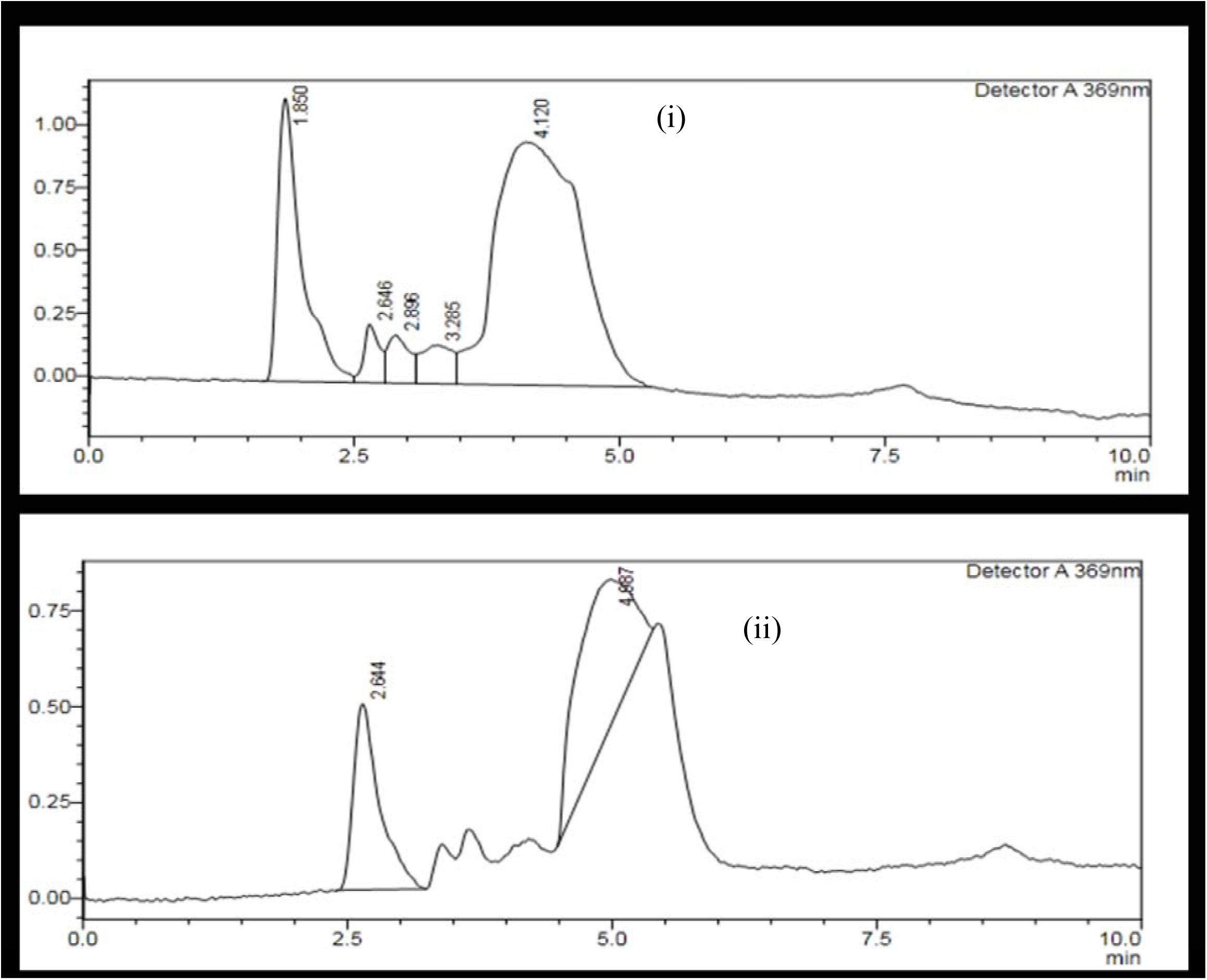
HPLC analysis of influent (i) and effluent (ii) of SBR operated with 1000 mg/L of mixed dyes.

**Fig 9.**
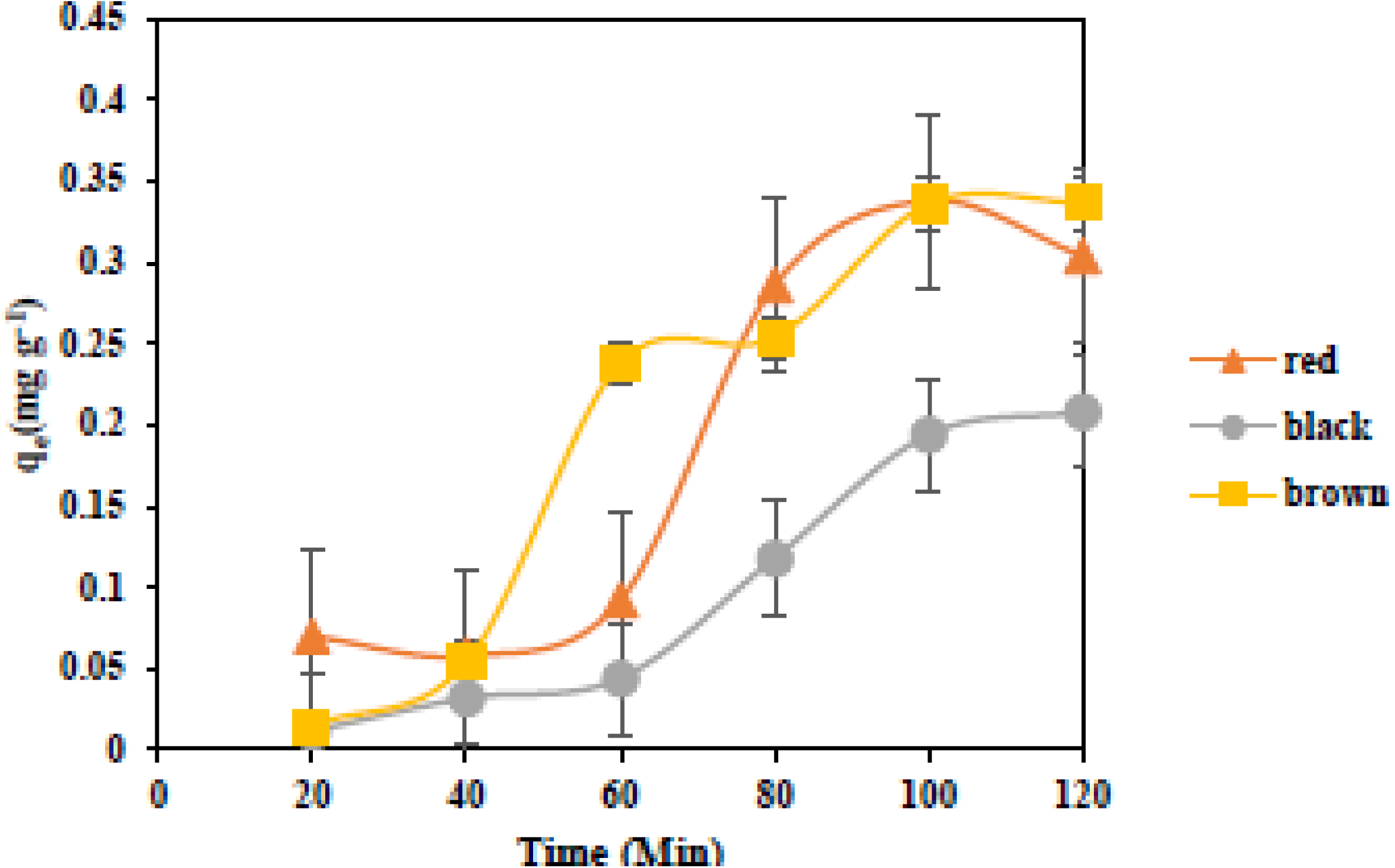
Biosorption of dyes in treated wastewater using DAS.

### Biosorption studies

Bioadsorption of dyes by the prepared adsorbent was observed in normal dye solution, this can be applied to the wastewater that was treated by the SBR and MBBR to enhance the decolorization percentage. Therefore to study the ability of dried activated sludge to adsorb the dyes present in treated wastewater was checked in 100 mg/L.

At 20 minutes the amount of RR adsorbed was 0.07 mg/g, 0.011mg/g of RB and 0.015mg/g of RBR. The adsorption gradually increased till 60 minutes after which there was a rapid adsorption noticed at 80 minutes. At 80 minutes, 0.286 mg/g, 0.118 mg/g and 0.254 mg/g of RR, RB and RBR was adsorbed and adsorption increased gradually till two hours after which the adsorption percentage decreased. At two hours, 0.304, 0.208, 0.337 mg/g of RR, RB and RBR was adsorbed. Adsorption after two hours showed no difference in decolorization. RBR and RR were adsorbed more by the biosorbent when tested with treated synthetic wastewater.

### Phytotoxicicty studies

Phytotoxicity studies using green gram showed 100 % germination in distilled water and treated wastewater. Untreated wastewater showed less amount of germination and it had reduced root and shoot length compared to control and treated wastewater in all three concentration i.e. 100, 500 and 1000 mg/L. This shows the amount of toxicity has been reduced in wastewater after treatment. Phytotoxicity was done for seven days. In 100 mg/L of treated wastewater and in the control 100 % of germination was observed. In untreated wastewater there was only 70 % of germination percentage. Since the dyes were not degraded in untreated wastewater, the chemicals and dyes inhibited the growth of the plant. There was 70 % of germination observed in the untreated wastewater. The root and shoot length of untreated wastewater was considerably small compared to that of control and treated wastewater proving that the wastewater has been treated and the dyes were degraded in treated wastewater. Since the dyes were not degraded in untreated wastewater, the chemicals and dyes inhibited the growth of the plant.

In 500 mg/L studies, the germination percentage for both control and treated wastewater was 100 %. Since the dye concentration is high in 500 mg/L only 60 % of germination was seen in untreated wastewater. In 1000 mg/L concentration, control and treated wastewater were able to aid in the plants’ growth. For both control and treated wastewater 100 % germination were observed whereas in the untreated wastewater only 60 % of germination was seen, this is because the dye concentration was very high in 1000 mg/L untreated wastewater and therefore it inhibited the growth of plants. The sprouts seeded in treated wastewater showed more root and shoot length compared to the control i.e. the root length was 3.96 cm and the shoot length were 18.28 cm, this may be due to the presence of activated sludge which acts as a fertilizer for the plants.

The phytotoxicity test with *Vigna radiata* showed that the high concentration of dyes can inhibit the growth of the plants. The results obtained were compatible to Karthikeyan and Kanchana, (2014). The growth of the plants in untreated wastewater was less compared to the growth of the plants in treated wastewater. In 500 mg/L studies, the treated wastewater plants showed more root and shoot length than the control, this may be due to the fertilizing quality of activated sludge used in the treatment of wastewater.

## Conclusion

Wastewater from textile industries has to be treated in order to avoid water pollution. If not treated, the aquatic life and human life will be affected due to the mutagenicity and toxic nature of the chemicals and dyes present in the textile effluent.

In the present study the synthetic wastewater containing mixed textile dyes were treated using a SBR. The treatment in SBR using activated sludge gave effective results. Further the studies can be continued as a comparative study by comparing the performance of SBR and a moving med biofilm reactor (MBBR) in different mixed dye concentration and by applying the decolorization process using the dried activated sludge by chemically activating and using it to post treat the wastewater that was treated in SBR. The sludge obtained after the treatment can be used further for remediation by using alkaliphilic bacterial consortium. Further shrubs and other phytoremediating plants can be used for completely removing the textile dye from the wastewater and soil.

## Acknowledgement

The Authors would like to acknowledge UGC-MRP-2017-19 for the Financial assistance to carry out the project.

---

**Tab 1.** Characteristics of the influent and effluent in SBR with wastewater containing 100 mg/L mixed dyes.

| **Parameters** | **Influent**<br>**(mg/L)** | **SBR effluent**<br>**(mg/L)** |
| --- | --- | --- |
| COD | 320 | 110 |
| BOD | 900 | 480 |
| Chloride | 600 | 400 |
| TDS | 400 | 350 |
| Hardness | 700 | 300 |

**Tab 2.**
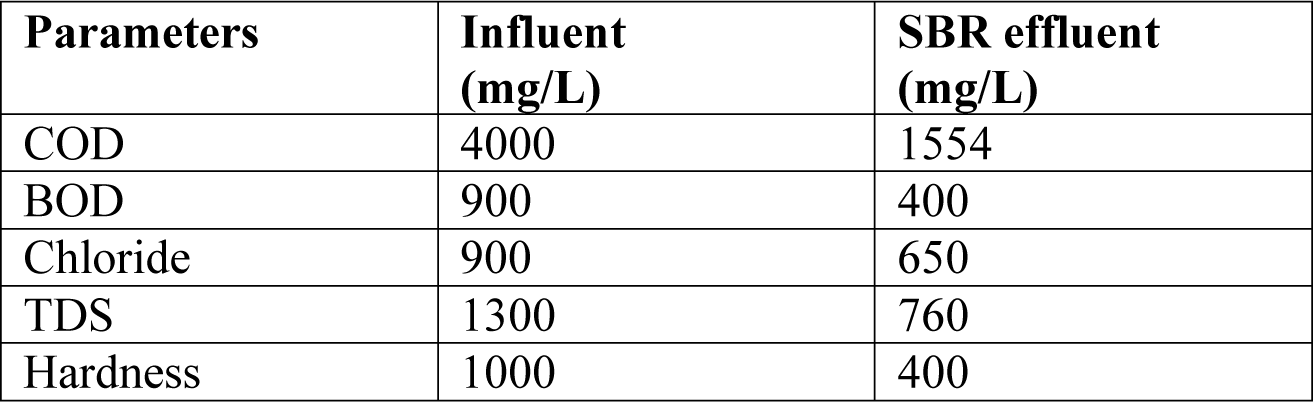
Characteristics of the influent and effluent in SBR with wastewater containing 500 mg/L mixed dyes.

**Tab 3.**
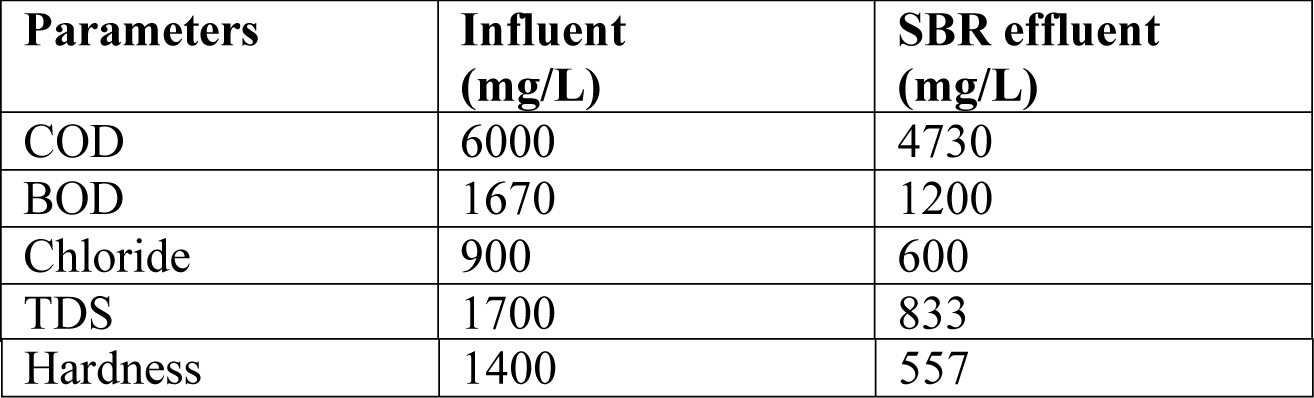
Characteristics of the influent and effluent in SBR with wastewater containing 1000 mg/L mixed dyes.

**Tab 4.**
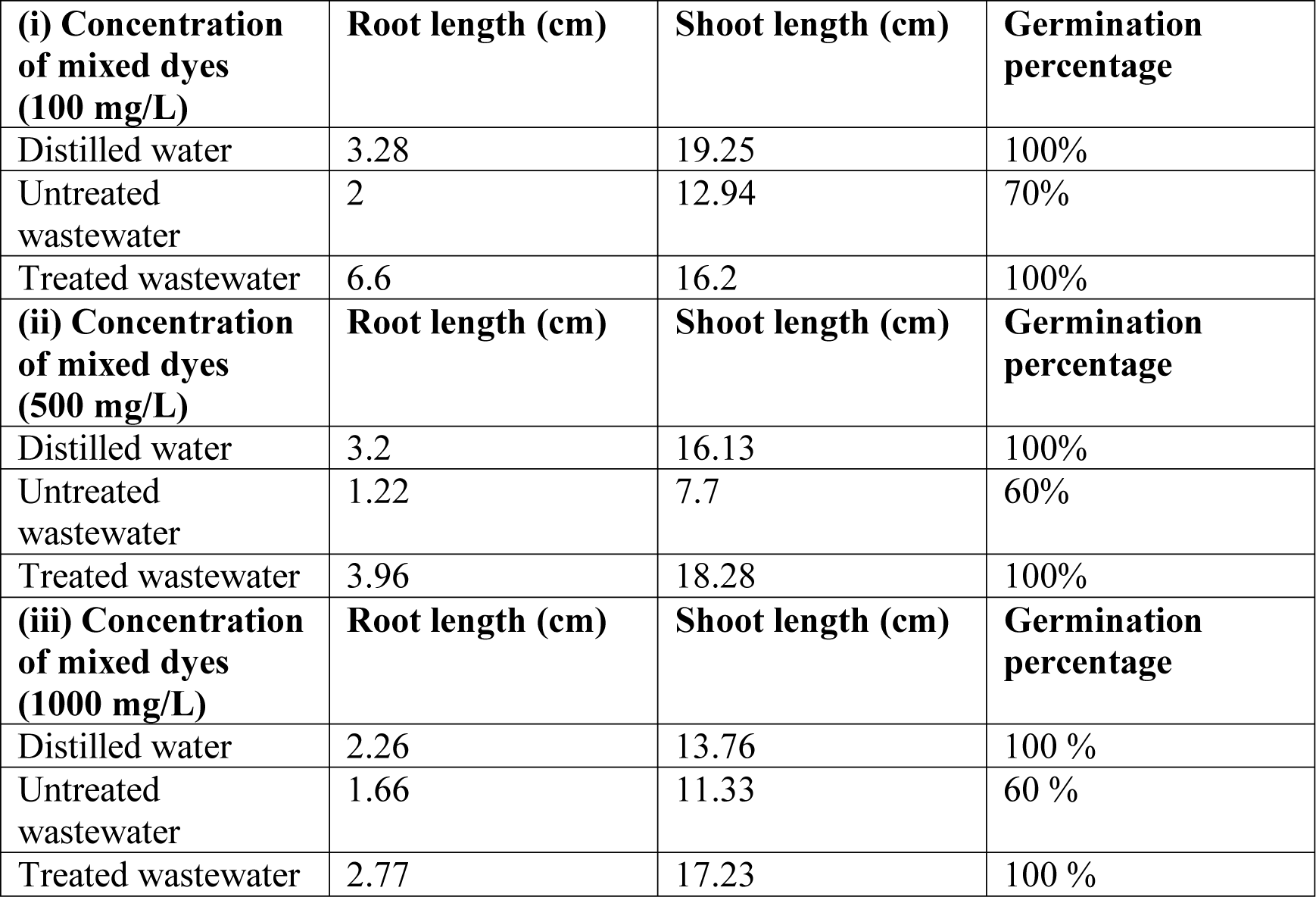
Phytotoxicity study using green gram with wastewater containing (i) 100 mg/L of mixed dyes (ii) 500 mg/L of mixed dyes and (iii) 1000 mg/L of mixed dyes.

